# Reproductive and ecological adaptations to climate underpin the evolution of sociality in lizards

**DOI:** 10.1101/2024.05.01.592000

**Authors:** B Halliwell, E. A O’Connor, T Uller, S Meiri, B.R Holland, C.K Cornwallis, G. M While

## Abstract

Identifying the environmental factors associated with group living is important for understanding how social systems originate, persist and diversify. In endothermic birds and mammals, living in social groups is associated with habitat constraints and harsh climatic conditions. We use phylogenetic comparative analyses to test whether climate and habitat have played similar roles in the evolution of social grouping in a globally distributed clade of ectothermic vertebrates, lizards (N_species_ = 1696). Social grouping was strongly associated with cool, dry climates across the lizard phylogeny. However, this climatic signature arose indirectly, by association with live birth (common in cool climates) and a reliance on rock crevices (common in dry climates), traits which increase parent-offspring associations and reduce offspring dispersal. In contrast, direct effects of cool temperature on the evolution of social grouping were marginal and restricted to live bearing species. Our results demonstrate that relationships between climate and sociality may result from climatic adaptations that go on to promote the emergence of grouping behaviour.

## Introduction

Evolutionary transitions from solitary to group living are strongly shaped by the environment. Climatic factors mediate the availability of resources, affecting the frequency, costs, and benefits of interactions among individuals (Wynne-Edwards, 1998; AlRashidi et al., 2010, 2011; Vincze et al., 2017; Lejeune et al., 2019). Other aspects of the environment, such as habitat structure and availability, can restrict dispersal and breeding opportunities, generating selection for increased social tolerance and cooperation (Emlen 1982; 1995; Komdeur 1992; Hatchwell & Komdeur 2000; Halliwell et al. 2017b). This has resulted in strong associations between environmental conditions and sociality in diverse clades across the tree of life (Jetz & Rubenstein 2011; Lukas & Clutton-Brock 2017; Cornwallis et al. 2017; Firman et al. 2020; Qi et al. 2023).

Most evidence that climate and habitat explain variation in social systems comes from complex forms of sociality, such as cooperative breeding (Rubenstein & Abbot 2017). Studies tend to focus on how the environment mediates the costs and benefits of independent breeding versus helping relatives to breed. For example, in birds and mammals, cooperative breeding is associated with climates where independent breeding is difficult, and the benefits of helping are high (Firman et al. 2020; Cornwallis et al. 2017; Lukas & Clutton-Brock 2017; Jetz & Rubenstein 2011). However, our understanding of the environmental drivers of sociality in vertebrates is potentially biased by a strong focus on endothermic species. We know comparatively little about the role of environment in ectothermic vertebrates, despite even greater potential for climate to mediate social behaviour (Moss & While 2021).

We use a global dataset of lizards to test how environmental factors have shaped the evolution of social grouping in this diverse clade of ectotherms. Environmental factors may have a more direct effect on social grouping in lizards compared to endotherms for several reasons. First, cool climates impose constraints on activity (Vidan et al. 2017), affecting dispersal and opportunities for social interactions (Price-Rees et al. 2014; Rat et al. 2020; Moss et al. 2023). Second, cool climates may alter kin structure by reducing polyandry (Olsson et al., 2011), and intraspecific aggression (Baird & May, 2003), with consequences for inclusive fitness.

Finally, climatic conditions may select for adaptations that affect opportunities for interacting with relatives. In particular, the evolution of live birth (viviparity) is associated with cool climates in lizards (Zimin et al. 2022) and appears to have facilitated transitions to social grouping, in part, by promoting parent-offspring associations (Halliwell et al. 2017).

Conversely, some aspects of environmental variation may have more general effects that promote sociality in both endotherms and ectotherms. Like birds and mammals, lizards require specific habitat features that can be limited or clustered across the landscape, often dependent on local climatic conditions. Such clustering can favour the evolution of social tolerance, especially if constraints on dispersal promote interactions between kin (Halliwell et al. 2017b; Le Galliard et al. 2003; Massot et al. 2002). However, whether such general environmental conditions can explain the evolution of sociality across diverse taxa remains unclear.

We integrated data on social grouping (Halliwell et al. 2017), breeding climates (Caetano et al. 2022; Harris et al. 2020), habitat preferences (Meiri 2024) and reproductive mode (Zimin et al. 2022) across 1696 lizard species. Sociality in lizards ranges from transient associations to stable transgenerational aggregations (Whiting & While 2017; While et al. 2019), but commonly occurs as family units based on delayed dispersal and prolonged parent–offspring associations (Whiting & While 2017). These simple social groups have evolved multiple times independently (Halliwell et al. 2017), making it possible to estimate how social grouping, climate and habitat preferences coevolve. We used phylogenetic comparative methods to 1) test if social grouping is explained by climate and habitat after accounting for viviparity; 2) evaluate alternative causal models for the relationships between these traits and; 3) test for differences in reconstructed trait values between nodes estimated to represent different stages in the evolution of social grouping.

## Methods

### Data Collection

#### Social grouping and reproductive mode

We used data on lizard social grouping from Halliwell et al. (2017) updated with reports of social grouping published between 2017 and 2024. Social grouping was assigned based on reports of social aggregations containing both adults and juveniles. One key challenge to such studies is assigning the absence of a trait. Species were classified as non-social if they met any of the following four criteria (see Halliwell et al. 2017 for details): (1) Other forms of parental care (e.g. nest construction, egg guarding) have been reported for the species with no mention of associations between adults and juveniles in any of the literature accessed; (2) Studies of life history, reproductive biology, or ecology involving active field sampling are available for the species, but do not report associations between adults and juveniles; (3) The species is well studied (citation count of ≥100 peer-reviewed publications according to Web of Science), but no reports of associations between adults and juveniles were found during the literature search; (4) complete absence of reports of social grouping for that taxonomic family.

Data on reproductive mode were extracted from Zimin et al., (2022). The final data set used for analyses contained 1696 species.

#### Climatic niche

The climatic niche of each species was estimated using data from the CRU TS v. 4.07 climate grids. This contains monthly temperature and precipitation estimates for 0.5° grid cells across all global land areas (excluding Antarctica) for the period between 1901 and 2022 (Harris et al. 2020). Species distribution maps from Caetano et al. (2022; also see Roll et al. 2017) were intersected with the CRU database to calculate the median value of mean monthly temperature (degrees Celsius) and precipitation rate (mm/month) across grid cells for each species. This resulted in a median value for each climatic variable for each species for each month of every year. From this, we only used data for the 6 months of the year that are most relevant to embryo development and early post-natal life in lizards, i.e., breeding season (April – September for N hemisphere; October – March for S hemisphere). From these data, we calculated a mean for each species for each climatic variable. These temperature and precipitation values (henceforth temperature and precipitation) represent averages over each species geographic range during the breeding season. Predictability in temperature and precipitation were measured via Colwell’s P (Colwell et al. 1974), an index that captures among-year variation in onset, intensity and duration of periodic phenomena ranging from 0 (completely unpredictable) to 1 (completely predictable). We chose Colwell’s P over measures of variability such as standard deviations because these metrics were less strongly correlated to temperature and precipitation in our data (Figure S1), and facilitate comparisons with other studies (Cornwallis et al. 2017; Fristoe et al. 2017; Griesser et al. 2017; Firman et al. 2020; Martin et al. 2020; Diamant et al. 2021). However, it is worth noting that the interpretation of predictability should be guided by an assessment of the relative contribution of constancy and periodicity to predictability estimates (Liu et al. 2024). These supporting analyses are provided as supplementary information (Figure S2). Temperature (after conversion to K) and precipitation values were log-transformed before calculating species means and Colwell’s P values.

#### Habitat associations

We extracted data on habitat associations from SquamBase (Meiri 2024). Habitat associations included 4 broad non-exclusive habitat types: arboreal (living and foraging on vegetation – 766 spp.), fossorial (henceforth “burrowing”, digging or foraging underground or under objects – 127 spp.), saxicolous (henceforth “rock dwelling”, living and foraging in and on rocks – 334 spp.) and terrestrial (moving and foraging on the ground and leaf litter – 990 spp.) coded as binary variables (i.e., is the species found in this habitat type? yes/no).

### Statistical Analyses

We used three complementary approaches to investigate coevolution between climate, habitat, reproductive mode and social grouping in lizards. First, we used phylogenetic mixed models to: 1) estimate regression coefficients between environmental predictors and social grouping while accounting for phylogenetic dependence; 2) explore interactions between environmental variables and viviparity on the probability of social grouping; 3) estimate phylogenetic correlations between environmental variables and social grouping using multi-response models. Second, we used phylogenetic path analyses to gain insight into the causal network underlying trait correlations. Third, we performed ancestral state reconstructions to test for differences in climatic niche and habitat associations among ancestral species (internal nodes) that did, and did not, give rise to sociality in descendant lineages. The R code necessary to replicate all analyses is included as supplementary information.

#### Testing for trait relationships while controlling for phylogeny

We used Bayesian phylogenetic mixed models (PMM) (MCMCglmm package; Hadfield 2010) to fit phylogenetic regressions that tested for associations between social grouping, reproductive mode, climatic niche and habitat associations while controlling for phylogenetic relatedness among species. In PMMs, phylogenetic random effects are modelled via (co)variance matrices that assume a Brownian motion model of evolution (Hadfield and Nakagawa 2010). We modelled social grouping as a binary trait (yes vs no) on the probit link scale using a “threshold” error distribution and each climate variable and habitat association as a fixed effect predictor. To test whether the effects of climate and habitat were independent of reproductive mode, we re-fit the model with viviparity (yes vs no) as an additional predictor and checked for changes in the significance of fixed effects.

Based on initial analyses identifying viviparity and rock dwelling as key predictors of social grouping (see results; Table S2) we fitted a series of PMMs to test whether these traits interact with climate variables (or each other) to influence the probability of social grouping. Specifically, a series of candidate models specifying all possible two-way interactions between both viviparity and rock dwelling, and each climate variable, as well as all three-way interactions between viviparity, rock dwelling, and each climate variable (Table S3). We then compared these models to a null model including only the main effects of viviparity and rock dwelling using deviance information criterion (DIC) as a measure of predictive performance that accounts for the effective degrees of freedom (EDF) in models including both fixed and (phylogenetic) random effects (Hadfield 2010). Although model scores such as WAIC or LOO may be preferable to DIC due to their asymptotic properties (Gelman et al. 2014), these options are not available for phylogenetically structured data in the Gibbs sampling framework of MCMCglmm.

#### Testing for phylogenetic correlations

Phylogenetic regressions model dependence in the residual error structure attributable to phylogeny, and therefore ‘control for phylogeny’ when estimating regression coefficients. However, when predictor variables also contain phylogenetic signal, multi-response (MR) PMMs offer a more informative decomposition of trait covariances (Westoby et al. 2023; Halliwell et al. 2022). We used MR-PMM to examine phylogenetic correlations between social grouping, viviparity, rock dwelling, burrowing, and each of the 4 climate variables (8 response variables in total). We excluded terrestrial and arboreal habitat associations as response variables in MR-PMM to minimise the number of estimated model parameters, as these habitat associations showed a weaker relationship with social grouping (social grouping occurs in only 4% of terrestrial species and 2.3% of arboreal species) compared with rock dwelling (11.3%) and burrowing (9.4%) (Table S4). We modelled the binary traits of social grouping, viviparity, rock dwelling and burrowing on the probit link scale using “threshold” error distributions. We modelled the continuous traits of (log) temperature, (log) precipitation, temperature predictability and precipitation predictability using a Gaussian error distribution. All continuous traits were z-transformed prior to analyses.

For the MR-PMM, we removed the global intercept to allow separate intercepts for each trait and specified an 8x8 unstructured phylogenetic covariance matrix for the random effects. The variance partitioning achieved with MR-PMM also allows covariances between traits on the residual level (i.e., covariances independent of phylogeny) to be estimated. However, for binary traits, the residual variance is not identifiable and is fixed in the prior specification of the model (Hadfield 2010). We therefore specified structured (diagonal) residual covariance matrices for the residual errors in MR-PMM and relied on phylogenetic regression to estimate trait relationships residual to phylogeny. For both PMMs and MR-PMMs, we incorporated phylogenetic uncertainty by fitting models over a sample of 100 candidate topologies from Tonini et al. (2016) and concatenated MCMC chains of parameters across fits for inference. Each tree was pruned to contain only those species for which data on reproductive mode, environmental variables and social grouping was available (n = 1696).

#### Testing for phylogenetic correlations separately for oviparous and viviparous species

We investigated whether correlations between social grouping, rock dwelling and temperature differed between oviparous and viviparous species by fitting a MR-PMM that estimated separate phylogenetic correlations between these traits for the two reproductive modes using at.level coding in MCMCglmm (see code in supplementary information). This approach doubles the number of estimated model parameters, asking more of the data (especially for viviparous species, n = 197). Therefore, we did not attempt it for the full MR-PMM including 8 response traits.

#### Model settings, diagnostics and parameter estimation

The prior settings used for each analysis are specified in the R code in the supplementary information. Following de Villemereuil (2018), we used parameter expanded priors for random effects in binary response models. The residual variance of binary traits was fixed to 1. Otherwise, an inverse-Wishart prior was specified for residual variances. Model parameter estimates are reported as posterior modes and 95% credible intervals.

We examined the convergence of MR-PMMs by fitting two separate MCMC chains and: 1) visually inspecting traces of the MCMC posterior estimates and their overlap; and 2) calculating potential scale reduction factors 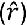, a convergence diagnostic test that compares within- and between-chain variance (Vehtari et al. 2019). 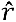values greater than 1.1 indicate chains with poor convergence properties. We confirmed 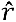 values were less than 1.1 for all parameters, though for most parameters 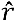was considerably lower 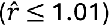.

For Gaussian traits, we estimated the proportion of variance explained by phylogenetic dependence (sometimes referred to as phylogenetic signal, λ) from the MR-PMM as, 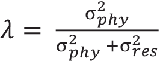, where 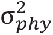and 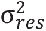are the phylogenetic and residual variances for a given trait from the fitted model, respectively. For binary traits, the residual variance was fixed (see above) and therefore we used the intraclass correlation coefficient to estimate phylogenetic signal. This is calculated as, 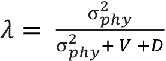, where V is the level of residual variance (additive over-dispersion) fixed in the prior specification (here, V = 1) and D = 1 is the distribution specific variance term for the binomial distribution with a probit link (Nakagawa & Schielzeth 2013).

We calculated the phylogenetic correlation between two traits, as:

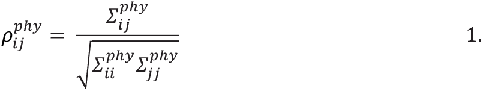

Where 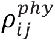is the phylogenetic correlation between traits *i* and *j*, 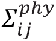is the phylogenetic covariance between traits *i* and *j*, and 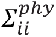is the phylogenetic variance in trait *i*. To evaluate evidence for conditional dependencies between traits, (i.e., relationships after controlling for covariances with other response traits) we also calculated partial phylogenetic correlations. To compute partial correlations, we substituted elements of the inverse trait-level phylogenetic covariance matrix, *∑*^*phy−*1^, into equation 1 following Halliwell et al. (2022) (also see Popovic et al. 2019). To evaluate the significance of correlation estimates, we checked whether the 95% HPD of the posterior distribution of parameter estimates crossed zero.

#### Phylogenetic Path Analysis

To evaluate different hypotheses of the causal network underpinning trait relationships, we conducted phylogenetic path analyses in the ‘phylopath’ package (van der Bijl 2018). We utilised frequentist approaches for these analyses due to the computational constraints on fitting many alternative causal models across a posterior sample of trees in a fully Bayesian framework. Specifically, based on results from phylogenetic mixed models (Figure 3a; Table S2), we defined 27 candidate models for the network of relationships between social grouping, viviparity, rock dwelling, precipitation, and temperature (Figure S3). We then fit this set of candidate models via maximum likelihood to each of 100 candidate topologies and summarised the distribution of point estimates generated for each path coefficient. We also examined the distribution of model weights (Figure S4) to assess the influence of phylogenetic uncertainty on model selection, and the distribution of standardized regression coefficients to assess the significance of trait relationships. Inferences were drawn by averaging over all candidate models, weighted by their relative evidence, with confidence intervals for regression coefficients calculated using 100 bootstrap replicates.

#### Ancestral State Reconstruction

We reconstructed ancestral states for reproductive mode and social grouping using hidden Markov models (HMM) in the R package corHMM (Beaulieu et al 2013; Beaulieu & O’Meara 2014). This modelling framework uses “hidden” states to parametrize multiple rate matrices, allowing transition rates in a discrete character to vary across the phylogeny (i.e., allowing transition rates to vary between clades). We chose this approach based on recent research indicating that the likelihood of evolving viviparity varies considerably across squamates, which has important effects on ancestral state reconstructions (Pettersen 2023; also see King & Lee 2015; Wright et al. 2015).

For each trait (e.g. reproductive mode), we fit HMM that allowed between 1 and 3 different rate categories from one state (oviparity) to the other (viviparity) across the tree. Reversals from viviparity to oviparity were prohibited because such events are thought to have been rare in the squamates (Blackburn 2015; Shine 2015; Griffith et al. 2015). We used AICc to compare models and identify the most likely number of rate categories (1-3) across a sample of 100 candidate topologies. A model with 2 rate categories was preferred for social grouping (lowest AICc for 92/100 trees) and a model with 3 rate categories was preferred for reproductive mode (lowest AICc for 74/100 trees). The preferred model for each trait was used in all subsequent analyses.

Joint estimates of ancestral states at internal nodes from HMMs were then used to identify transitions in social grouping for oviparous and viviparous species separately, by classifying internal nodes according to their estimated state and that of their descendants, e.g., oviparous non-social with at least one social descendant (O.NSG_O.SG). Nodes inferred to have undergone simultaneous transitions in reproductive mode and social grouping (e.g., O.NSG_V.SG) were treated as NA. These unresolved nodes mean we are likely to underestimate the number of transitions to social grouping within our sample, especially for viviparous species (Table S5), which may influence our results. Therefore, we conducted sensitivity analyses in which these nodes were included, and confirmed equivalent results whether simultaneous transitions were treated as reproductive mode or social grouping (Figure S5) changing first in the sequence (e.g., O.NSG_V.SG treated as V.NSG_V.SG or O.SG_V.SG).

Following Cornwallis et al. (2017), we used these node classifications to test for differences in reconstructed trait values between nodes representing different stages in the evolutionary history of social grouping. For temperature, we used a PMM with temperature as the only response trait, node transition classification (e.g., O.NSG_O.SG) as a fixed effect, and a phylogenetic random effect linked to internal nodes. This model uses observed temperature values from species at the tips to reconstruct values at internal nodes and estimate fixed effects for each node classification category. Using the posterior distribution of fixed effects estimated for each node classification category, we constructed contrasts that test different hypotheses about the sequence of evolutionary events. Specifically, to test if temperature values differed between the ancestors of social and non-social lineages (i.e., to test for differences in temperature at the origin of social grouping), we compared reconstructed values for non-social ancestors that gave rise to only non-social descendants with those of non-social ancestors that gave rise to at least one social descendants (e.g., O.NSG_O.NSG - O.NSG_O.SG). To test for changes in temperature values following the emergence of social grouping, we compared non-social ancestors that gave rise to at least one social descendant with social ancestors that gave rise only to social descendants (e.g., O.NSG_O.SG - O.SG_O.SG). To test for differences in temperature values associated with the maintenance of social grouping, we compared non-social ancestors that gave rise only to non-social descendants with social ancestors that gave rise only to social descendants (e.g., O.NSG_O.NSG - O.SG_O.SG). The same procedure was repeated for ancestral state reconstructions of rock dwelling.

For each comparison, we evaluated significance by subtracting the vector of posterior samples for the fixed effect of the first node category from those for the second node category (e.g.,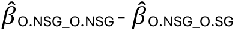) and checking whether the 95% credible interval of the resulting distribution overlapped zero (coloured bars in Figure 4). Finally, to test for differences between reproductive modes in the results of these contrasts, we subtracted relevant contrast distributions for viviparous species from those for oviparous species for each comparison (black bars in Figure 4).

## Results

The species included in our analysis occupied a wide range of climates (Figure 1), representing almost the entire global range of environments occupied by lizards (Pie et al. 2017; Roll et al. 2017). Social grouping occurred in 71 species (out of 1696, 4.2%) across 14 families, including 53 viviparous species (out of 197; 26.9%) and 18 oviparous species (out of 1499; 1.2%). Reconstructions from a hidden Markov model indicate that social grouping has arisen approximately 15 times in this sample of species (mean number of transitions (± 95% CI): oviparous lineages = 8.5 (5, 12); viviparous lineages = 6.1 (4, 11.5)) (Figure 2), though exclusion of uncertain nodes means the true number of independent origins is likely to be higher for viviparous species (see methods; Table S5). Based on the proportion of branch time spent in a state of oviparity (93.3%) verses viviparity (6.7%) from stochastic character mapping (Figure 2), these reconstructions indicate the transition rate to social grouping is at least 10 times faster from a background of viviparity.

**Figure 1.**
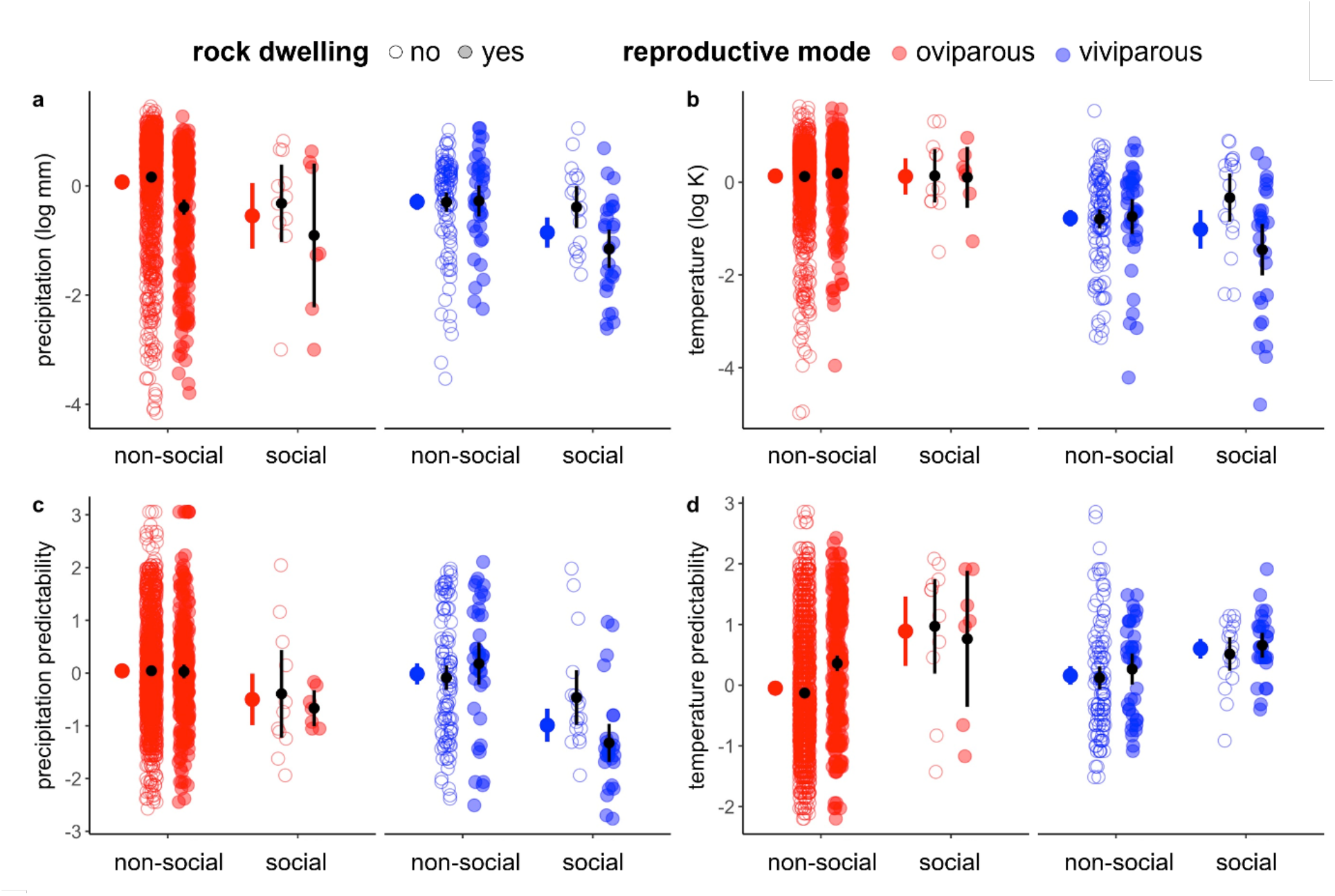
Reproductive (viviparity) and ecological (rock dwelling) adaptations to climate create a strong climatic signature of social grouping in lizards. Climate niche variables (z-scale) for lizard species (n = 1687) grouped by reproductive mode, rock dwelling and presence of social grouping. Means and 95% confidence intervals for each group are shown in black. Means and 95% confidence intervals for each combination of reproductive mode and social grouping (i.e., pooling across presence/absence of rock dwelling) shown to the left of each group combination. **a** log mean monthly precipitation during the breeding season. **b** log mean temperature during the breeding season. **c** inter-annual autocorrelation of mean monthly precipitation during the breeding season. **d** inter-annual autocorrelation of temperature during the breeding season. Outliers (n = 9) excluded for plotting.

**Figure 2.**
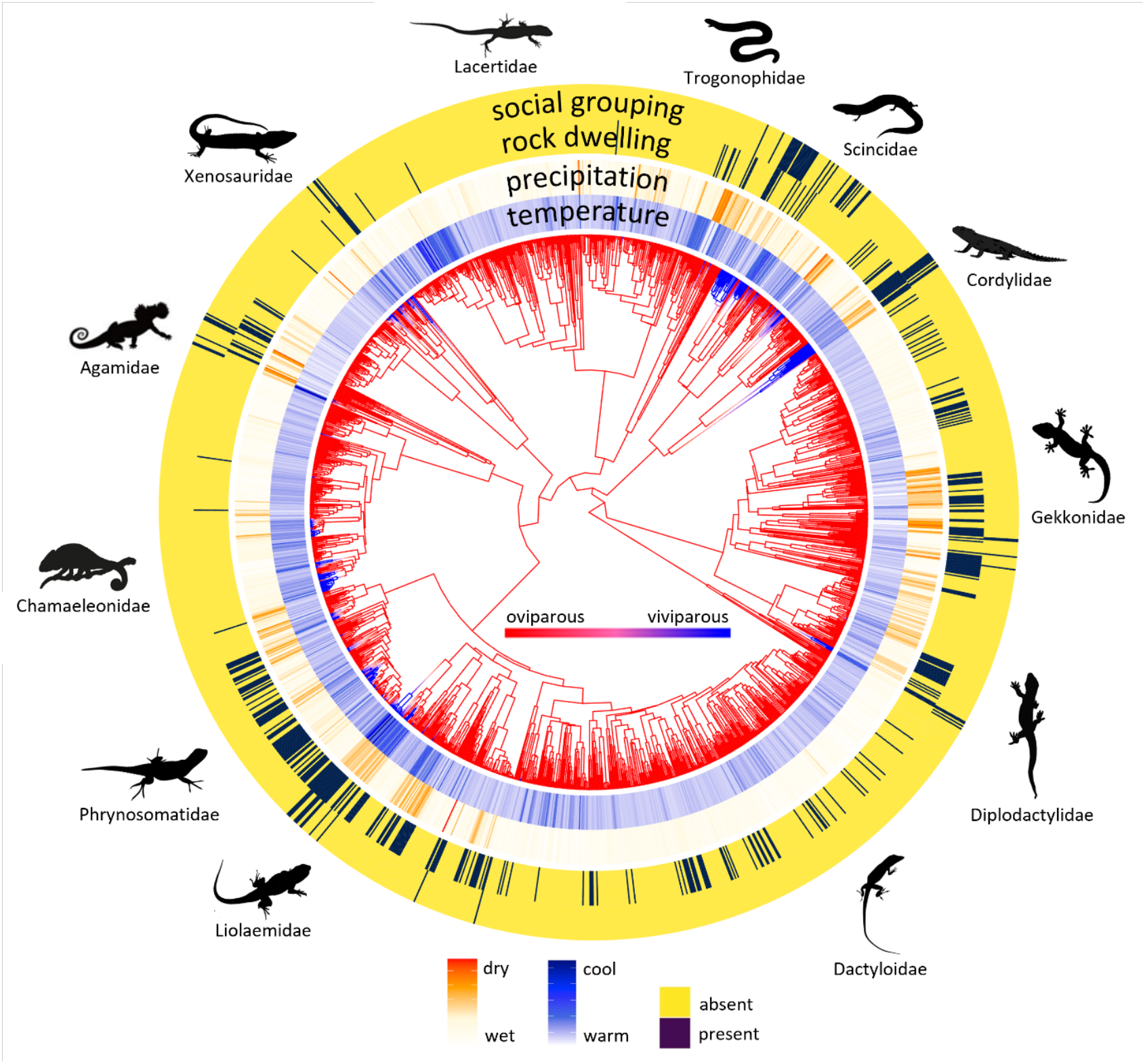
The phylogenetic distribution of viviparity, rock dwelling and climatic niche in lizards indicates that social grouping is a product of past adaptations and current ecological challenges. Branch colour represents the probability of oviparity (red) and viviparity (blue) based on 100 stochastic character histories simulated from a hidden Markov model with three rate classes (see Methods). Mean breeding-season temperature and precipitation (inner), as well as presence/absence of rock dwelling and social grouping (outer) among extant taxa (n = 1696) is indicated.

Social grouping was more likely to occur in species that occupied cool climates and rocky substrates (P_MCMC_ values from phylogenetic regression <0.001; Table S1). Despite trends in the data suggesting social species occur in drier climates with more predictable temperature and less predictable precipitation (Figure 1), main effects of all other climate and habitat predictors were non-significant (Table S1). One explanation for the relationship between temperature and social grouping is that species occupying cooler climates are more likely to be viviparous (Zimin et al. 2022), which in turn promotes sociality (Halliwell et al. 2017). In support of this hypothesis, the effect of cool climate disappeared when viviparity was included as a predictor in the model, while the positive effect of rock dwelling remained unchanged (Table S2).

There was a positive phylogenetic correlation between social grouping and rock dwelling, and a negative phylogenetic correlation between social grouping and temperature (Figure 3a; Table S7), indicating conserved relationships between these traits across the lizard phylogeny. However, consistent with our phylogenetic regression analyses, there was no evidence for a partial phylogenetic correlation between social grouping and temperature (i.e., when controlling for viviparity). In contrast, there were partial phylogenetic correlations between social grouping and both viviparity and rock dwelling (Figure 3a; Table S7). These results indicate independent coevolutionary relationships between each of these traits and social grouping that go beyond correlated responses to climate. Furthermore, a negative partial phylogenetic correlation between rock dwelling and precipitation (Figure S6; Table S7) suggests that, in addition to promoting transitions in reproductive mode, climate may also have indirect effects on the evolution of social grouping by influencing the probability that species occur on rocky substrates.

**Figure 3.**
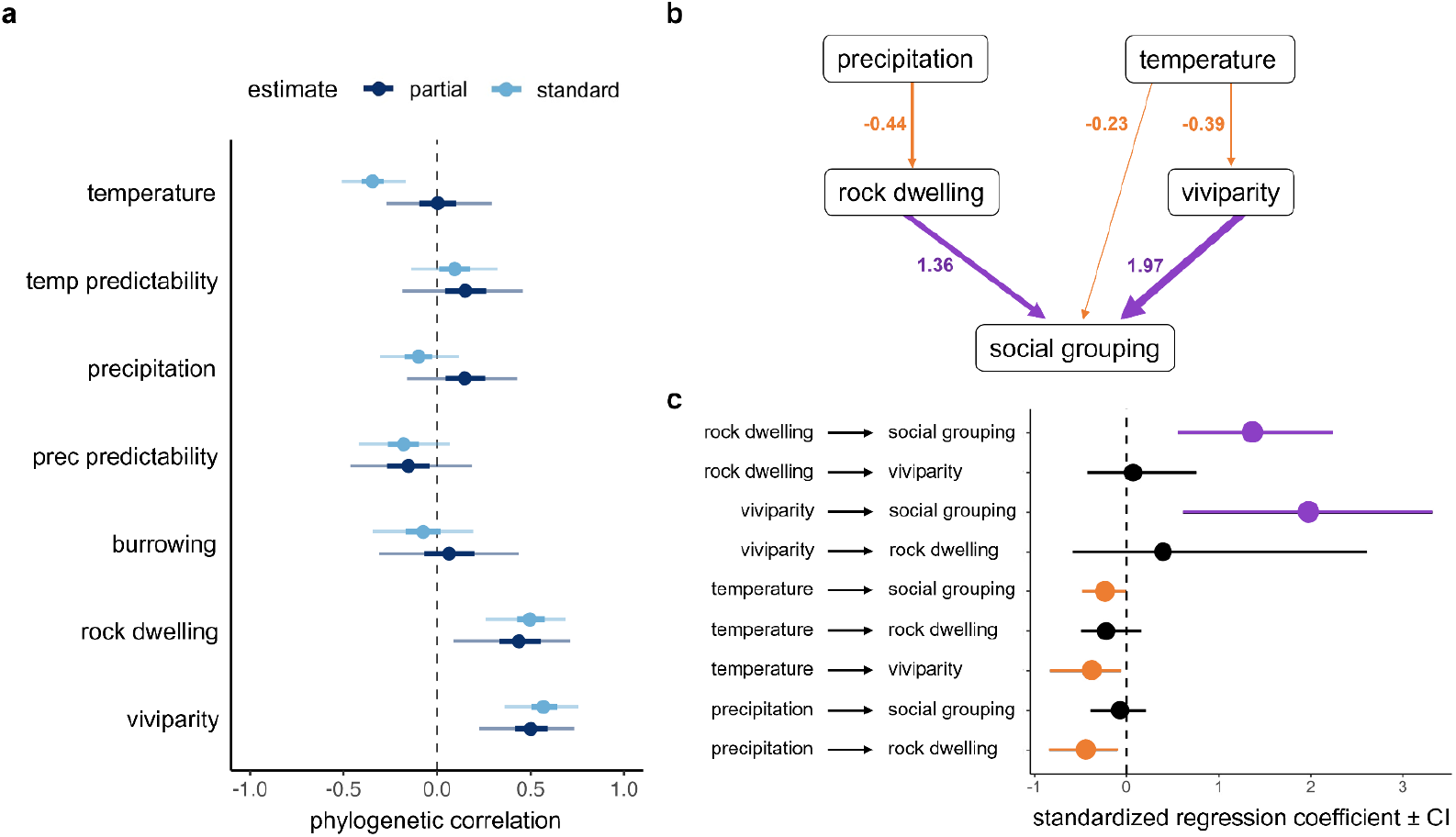
Viviparity and rock dwelling independently promote the evolution of social grouping across the lizard phylogeny, with limited evidence for direct climatic effects. **a** Phylogenetic correlations between species traits and social grouping from a MR-PMM fit over 100 candidate topologies. Estimates in light and dark blue represent standard and partial correlation coefficients. Partial correlations represent the relationship between two response variables after controlling for variation explained by other response variables. Points represent the posterior means, with heavy and light whiskers indicating the 0.5 and 0.95 credible intervals for each estimate. All pairwise correlations shown in Table S7. **b** Weighted average model from phylogenetic path analysis. Significant positive and negative paths shown in purple and orange, respectively. Coefficients associated with paths represent the mean standardized regression coefficient for each path across 100 candidate topologies. **c** Summaries of standardized regression coefficient estimates from weighted average model. Points and confidence intervals represent means and 95% confidence intervals for each coefficient estimate based on 100 bootstrap replicates across each of 100 candidate topologies.

There was limited evidence for social grouping being influenced by interactions between climate, viviparity and rock-dwelling (Table S3). The preferred model (lowest DIC for 87/100 trees) contained an interaction between viviparity and rock-dwelling, indicating that live-bearing species in rocky habitats were more likely to be social, but this effect was weak (P_MCMC_ = 0.07; Figure S7). Furthermore, this model was significantly preferred (δDIC > 2) over a null model containing just the main effects of viviparity and rock dwelling for only 23/100 trees (mean δDIC = 1.47; Table S3).

Phylogenetic path analyses indicated a strong positive influence of viviparity and rock dwelling, and a weak negative influence of temperature, on social grouping (Figure 3b,c). The model obtained by averaging over all candidate models, weighted by their relative evidence, also indicated a negative influence of temperature on both viviparity and rock dwelling, and a negative influence of precipitation on rock dwelling (Figure 3b). Each of these effects were significant based on the distribution of standardized regression coefficients across trees in the weighted model (Figure 3c).

Based on strong phylogenetic signal in all traits and environmental variables (Table S6), we performed ancestral state reconstructions to assess the effects of cool climates and rock dwelling across three key evolutionary stages: the origin, following the emergence, and during the maintenance of social grouping. There was a higher probability of rock dwelling at the origin of social grouping in both oviparous (marginal) and viviparous species, consistent with a general causative role of rock dwelling in the initial emergence of social grouping behaviour (Figure 4). Marginally cooler temperatures were also found at the origin of social grouping, but only for viviparous species. Consistent with these findings, rock dwelling and social grouping showed partial phylogenetic correlations across both oviparous and viviparous species, where- as temperature was only marginally negatively correlated to social grouping across viviparous species (Figure S8; Table S8).

**Figure 4.**
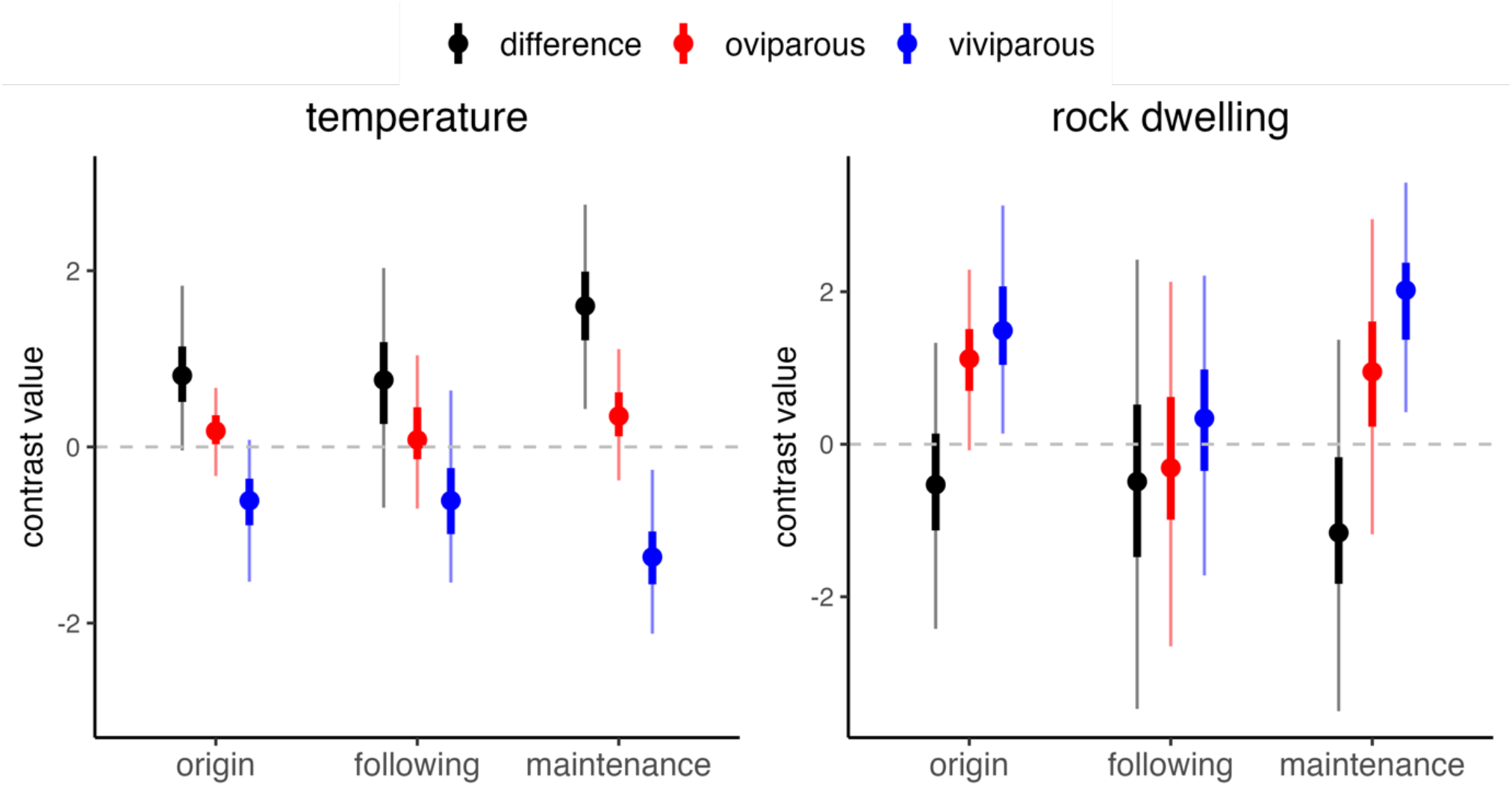
Rock dwelling promotes the emergence of social grouping regardless of reproductive mode, while effects of cool temperature are restricted to viviparous species. Contrasts between node transition categories representing different stages in the evolution of social grouping (see methods) for temperature (left) and rock dwelling (right). Values shown are posterior means (points) with 50% (heavy wicks) and 95% (light wicks) credible intervals calculated from phylogenetic mixed models fit across 100 candidate topologies. Contrasts for oviparous and viviparous lineages are displayed in red and blue, respectively. The difference between contrasts for oviparous and viviparous species are shown in black.

## Discussion

Climate has been repeatedly implicated as a causal factor mediating the evolution of vertebrate social behaviour (Jetz & Rubenstein 2011; Lukas & Clutton-Brock 2017; Firman et al. 2020). We show that social grouping in lizards also exhibits a strong climatic signature, typically occurring in species that occupy cool, dry climates. However, closer examination reveals that these effects are largely explained by association with viviparity and rock dwelling, both of which promote social grouping. Viviparity is an adaptation to cool climates in reptiles (Zimin et al. 2022; Ma et al. 2018) and is thought to promote group living by increasing the opportunity for interactions between parents and offspring following birth (Halliwell et al. 2017). Rock dwelling is strongly associated with dry climates but may also increase opportunities for social interaction among kin by promoting territoriality and ecological constraints on offspring dispersal (see below). Thus, associations between viviparity, rock dwelling and climate create a strong, yet largely indirect, climatic signature of social grouping.

Our finding that direct effects of climate on social grouping in lizards are weak contrasts with those from endothermic vertebrates (i.e., birds and mammals) where family life and cooperative breeding are often strongly associated with climate (Jetz & Rubenstein 2011; Cornwallis et al. 2017; Griesser et al. 2017; Firman et al. 2020; Qi et al. 2023), and harsh environments are thought to increase the relative benefits of social behaviour (Hatchwell 2000; Covas et al. 2008; Rubenstein et al. 2011). These differences can be understood by considering two important points.

First, associations between climate and complex social behaviour in birds and mammals are largely related to offspring provisioning (Qi et al. 2023; Cornwallis et al. 2017; Jetz & Rubenstein 2011). In this context, climate influences resource availability and thus the costs and benefits of prolonged investment in parental and alloparental care (Hatchwell 2009; Hatchwell & Komdeur 2000). This is likely tied to higher energy demands of endotherms which, all else being equal, make feeding offspring more important compared to ectotherms (Beekman et al. 2019; 2019b). There is no evidence that lizards provision their offspring and no evidence of alloparental care (While et al. 2014), potentially decoupling the functional links between climate, resource availability and social behaviour that underpin climatic signatures of social evolution in endothermic vertebrates. Conversely, climate may have distinct effects on the evolution of social behaviour in ectotherms due to inherent differences in thermal physiology (see introduction). In particular, thermal constraints on activity may promote kin sociality in lizards by stabilising social networks and reducing rates of extra-pair paternity (Moss & While 2021; Olsson et al 2011). We find some support for cooler temperatures promoting the emergence of social grouping, but only in viviparous species. This result suggests that direct effects of cooler temperatures may operate only when other conditions supporting the emergence of social grouping (i.e., viviparity) are already present. Living in generally cool climates may also mean that, for viviparous species, further reductions in temperature represent a direct metabolic constraint on activity, where-as in warmer regions (where oviparous species predominate) breeding-season temperatures rarely impose constraints on activity.

Second, our results indicate that climatic adaptations, more than climate per se, have mechanistic links to the evolution of social grouping in lizards. Climate often results in selection on a suite of adaptations which in turn select for social grouping. For example, climatic adaptation often involves morphological and life history traits (e.g., longevity, late maturation, and large body sizes), that can also mediate both the opportunity for kin groups to form, and the costs and benefits of grouping behaviour (e.g., Hatchwell & Komdeur 2000; Covas & Griesser 2007; Downing et al. 2015; 2020; Scharf et al. 2015; Griesser et al. 2017; Firman et al. 2020). Our analyses support this indirect association between climate and social grouping. Indeed, both reproductive (viviparity) and ecological (rock dwelling) traits were associated with climate adaptation and were shown to promote social grouping in lizards. Furthermore, we find marginal support for a positive interaction between viviparity and rock dwelling, suggesting these traits may act in concert to expedite the evolution of social grouping. This supports the observation that social grouping occurs most commonly in viviparous, rock dwelling clades (e.g., among the Australasian Scincidae, American Liolaemidae and African Cordylidae; Figure 2).

An additional novel finding of this study is that social grouping in lizards is associated with rocky habitats, and that occurrence in rocky habitats itself is associated with drier climates (Figure S6; Table S7). As ectotherms, lizards rely on refuge sites that offer specific microclimates and access to basking areas. Suitable refugia are often limiting, and habitat structure is particularly important in determining population dispersion and opportunities for offspring dispersal (e.g., Halliwell et al. 2017b; Botterill-James et al. 2016; Gardner et al 2016). As rock dwelling promotes social grouping in both oviparous and viviparous species, it follows that rocky habitats may have important effects on offspring dispersal regardless of reproductive mode. Reliance on persistent, defendable or manipulable home sites is consistent with the evolution of group living in other systems, such as transitions to eusociality in invertebrates (Howard & Thorne 2011; Nowak et al. 2010; Crespi 2001) and group living in rodents (Epsenberger 2001; Ebensperger & Hayes 2016). In contrast, we did not find an association between social grouping and fossoriality (burrow dwelling) as might be expected if general benefits related to shared refuge sites are an important driver of social grouping (e.g., Shah et al. 2003; Rabosky et al. 2012; also see Leu et al. 2011). However, the possibility to extend or excavate new borrow systems may mean that burrows are not limiting in the same sense as rock crevices.

In summary, we find the strong climatic signature of social grouping in lizards can be explained by association with viviparity and rock dwelling; reproductive and ecological adaptations to cool, dry climates that ultimately drive the emergence of grouping behaviour. Direct effects of climate are weak but suggest that cool temperatures may further promote the emergence of social grouping in viviparous lineages. Our results indicate that relationships between the environment and sociality are a product of interactions between past adaptations and current ecological challenges. Disentangling such effects can be difficult, requiring large datasets and sophisticated comparative methods. However, we show that such an approach is sometimes necessary to understand how and why social organisation varies across the tree of life.

## Acknowledgements

We thank Luke A. Yates for discussions and technical advice regarding Bayesian hierarchical modelling. BH was partially supported by the ARC Centre of Excellence for Plant Success in Nature and Agriculture (CE200100015). CKC was supported by the Knut and Alice Wallenberg Foundation (Wallenberg Academy fellowship 2018.0138).

