## Supplementary Figures for "Reproductive and ecological adaptations to climate underpin the evolution of sociality in lizards"

1. School of Natural Sciences, University of Tasmania, Australia
2. ARC Centre of Excellence for Plant Success in Nature and Agriculture, University of Tasmania, Australia
3. Department of Biology, Lund University, Lund, Sweden
4. School of Zoology, Faculty of Life Sciences, Tel Aviv University, Tel Aviv, Israel
5. The Steinhardt Museum of Natural History, Tel Aviv University, Tel Aviv, Israel

* Corresponding Author:

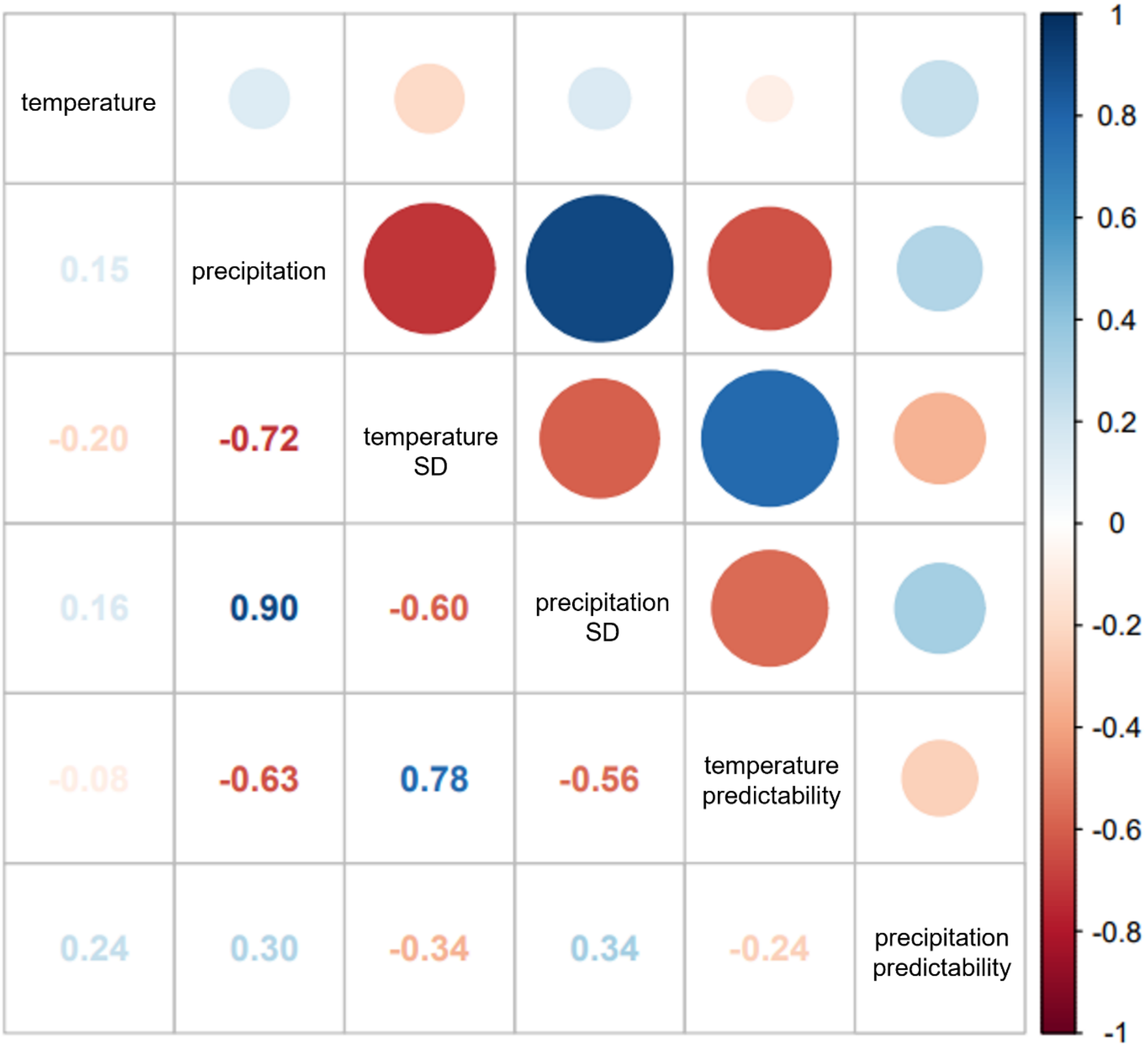


Figure S1. Correlations between means, standard deviations and predictability scores of breeding season temperature and precipitation for the lizard species included in analyses (N = 1696).

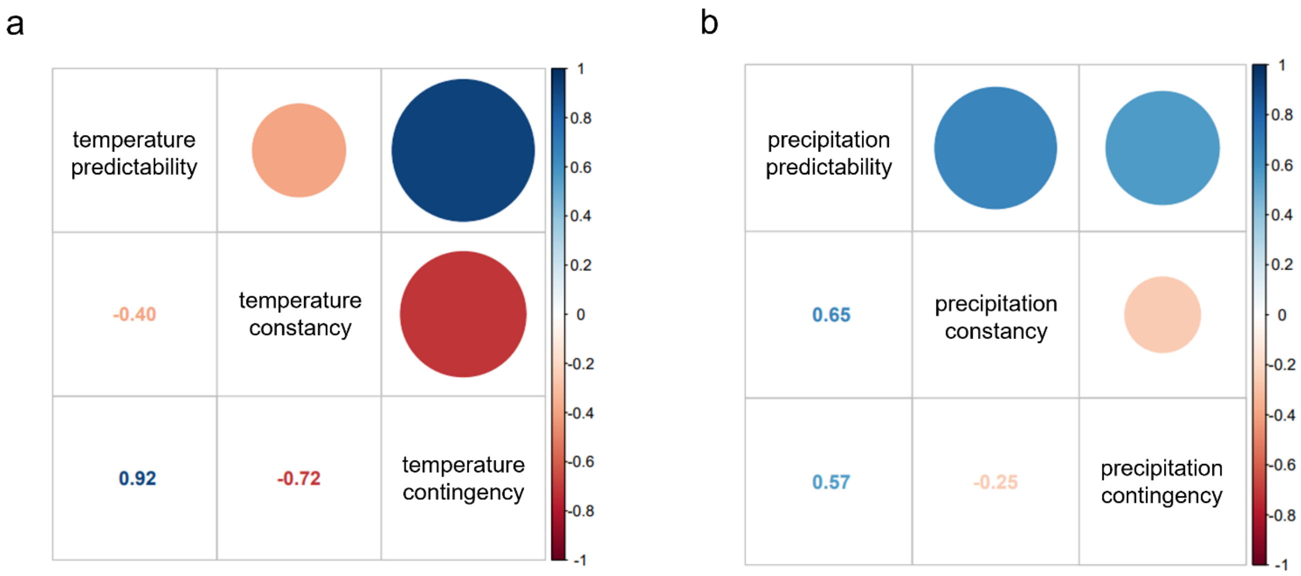


Figure S2. Correlations between predictability, constancy, and contingency scores for breeding season temperature (**a**) and precipitation (**b**) for the lizard species included in analyses (N = 1696).

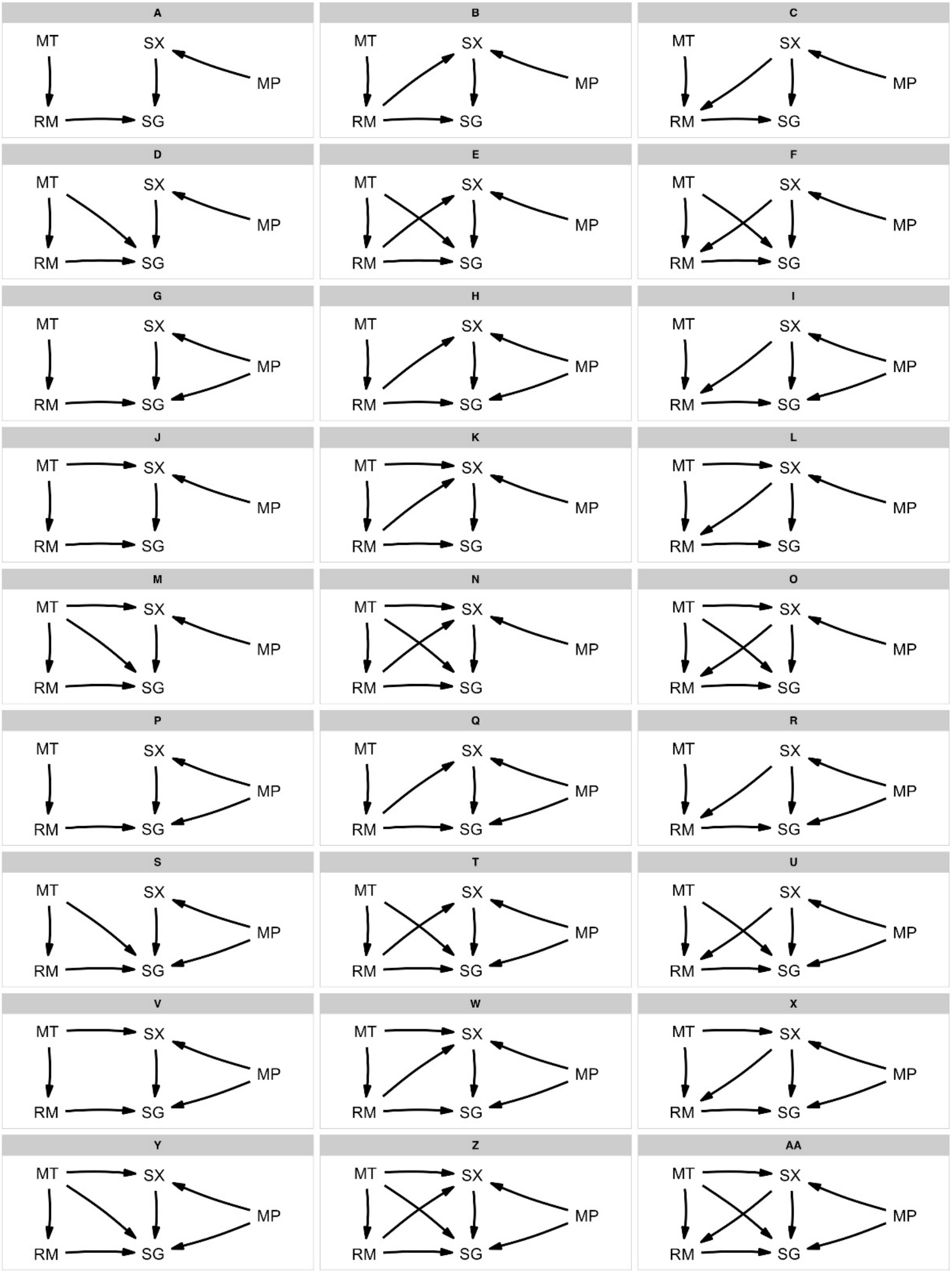


Figure S3. Candidate causal models between temperature (MT), precipitation (MP), viviparity (RM), rock-dwelling (SX) and social grouping (SG) evaluated by phylogenetic path analyses.

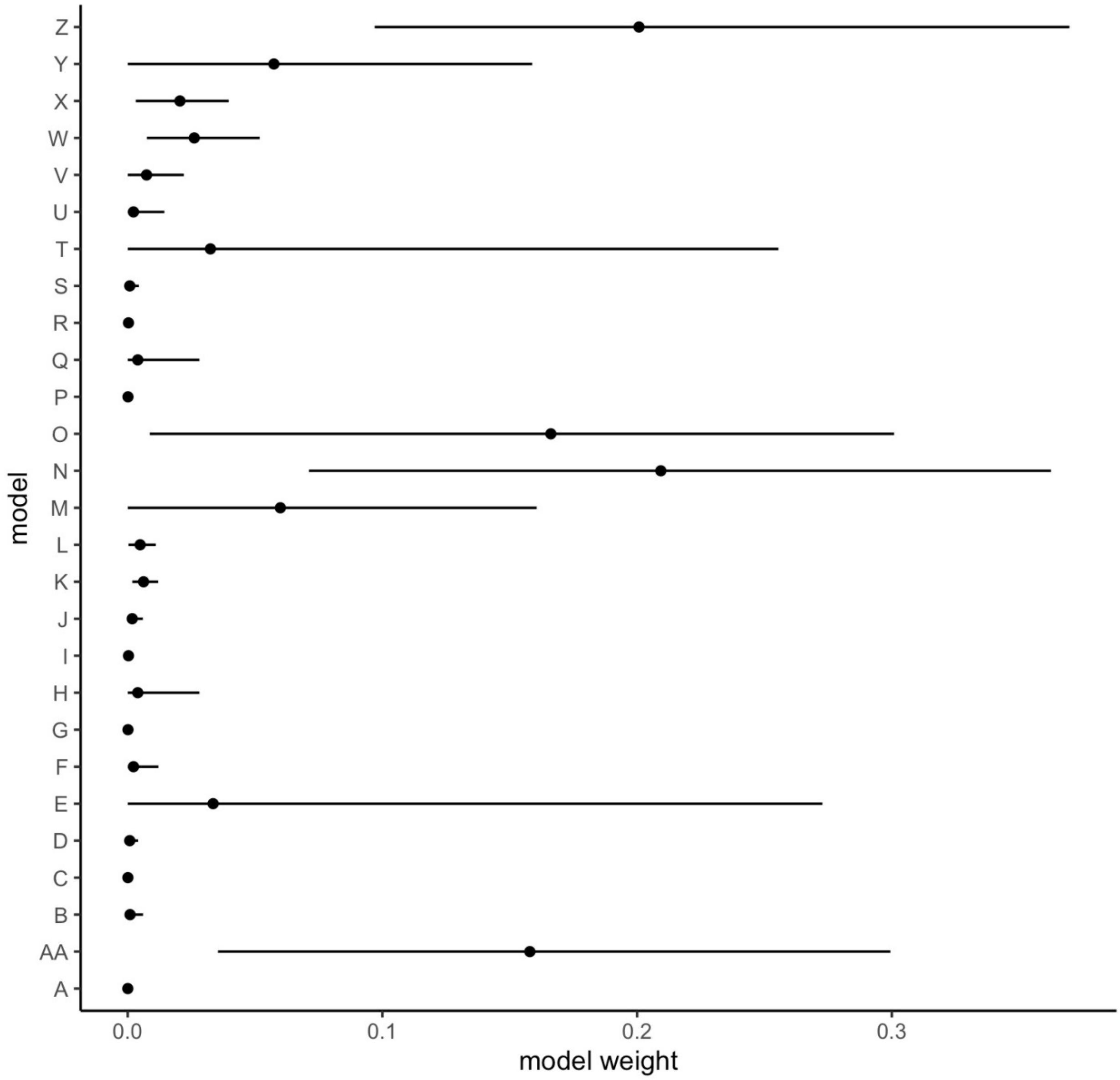


Figure S4. Model weights from phylogenetic path analyses. Points and whiskers show mean and 95% confidence intervals across 100 candidate topologies.

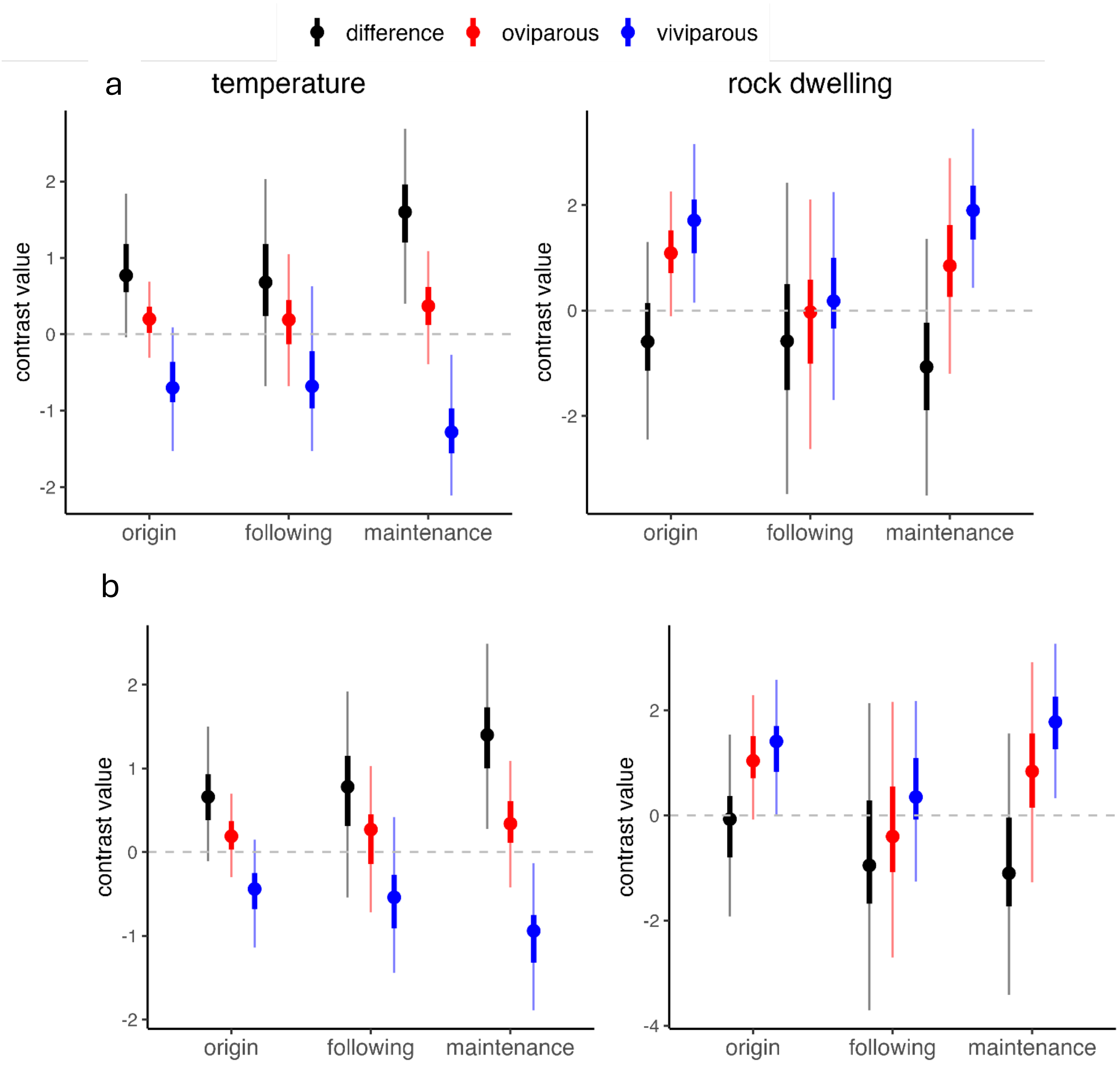


Figure S5. Contrasts between node transition categories representing different stages in the evolution of social grouping for temperature (left) and rock dwelling (right). (**a**) simultaneous transitions in social grouping and reproductive mode treated as reproductive mode changing first; (**b**) simultaneous transitions in social grouping and reproductive mode treated as social grouping changing first. Points represent posterior means, with heavy and light whiskers indicating the 0.5 and 0.95 credible intervals for each estimate.

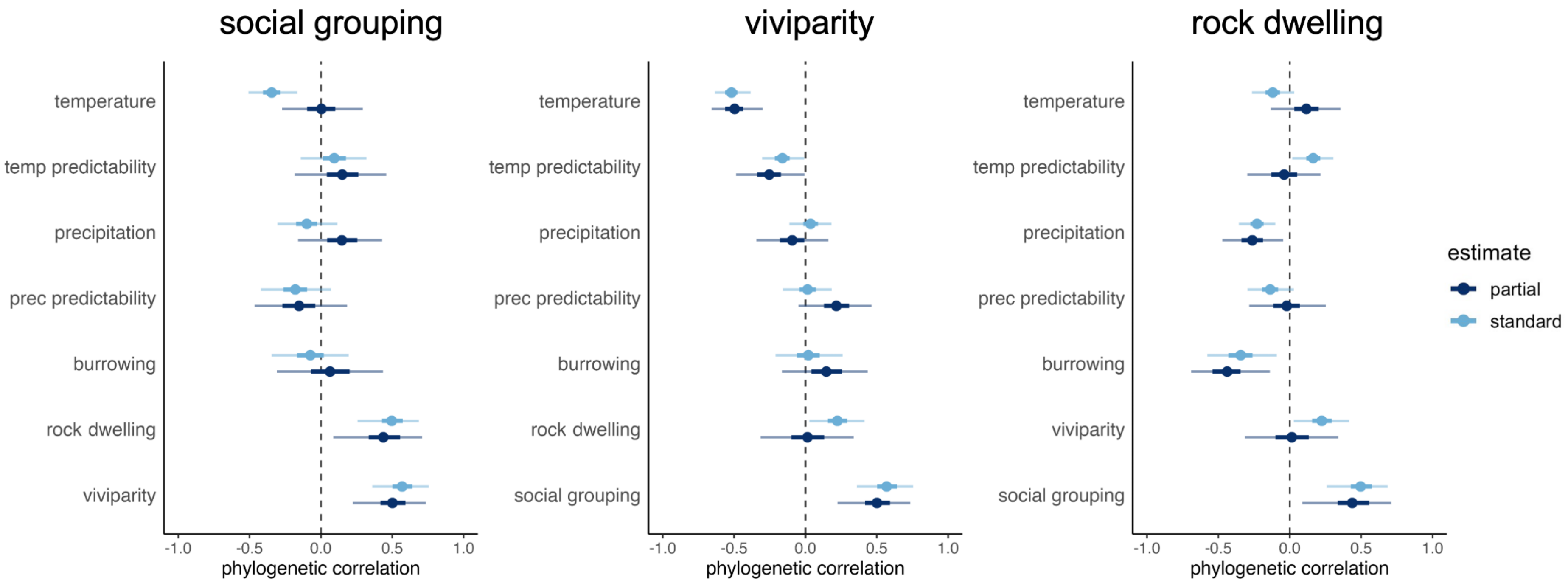


Figure S6. Phylogenetic correlations between species traits and a focal trait (title for each panel) from a MR-PMM fit over 100 candidate topologies. Estimates in light and dark blue represent standard and partial correlation coefficients, respectively. Partial correlations represent the relationship between two variables after controlling for variation explained by other predictors. Points represent the posterior means, with heavy and light whiskers indicating the 0.5 and 0.95 credible intervals for each estimate. Estimates for which the 0.95 credible interval does not cross zero are considered statistically significant. All pairwise correlations shown in Table S7.

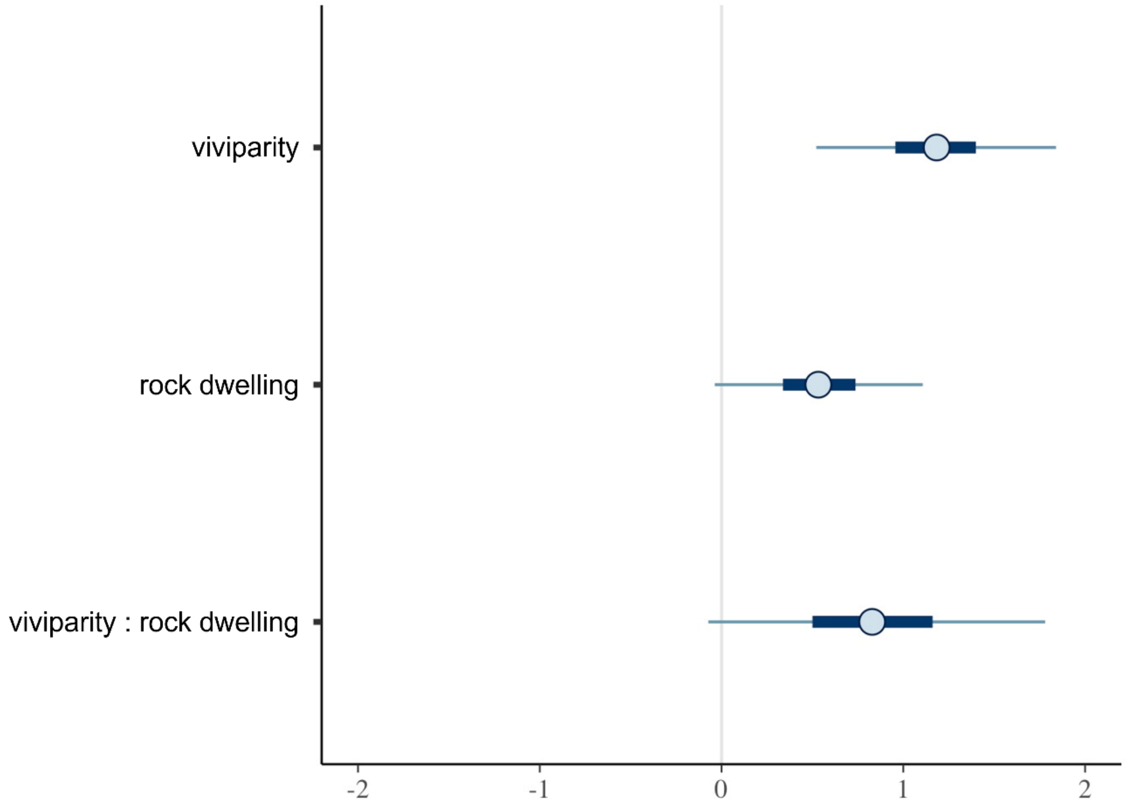


Figure S7. Coefficient estimates for fixed effects from the interaction model preferred by DIC (model 4 in table S3). Points represent posterior means, with heavy and light whiskers indicating the 0.5 and 0.95 credible intervals for each estimate.

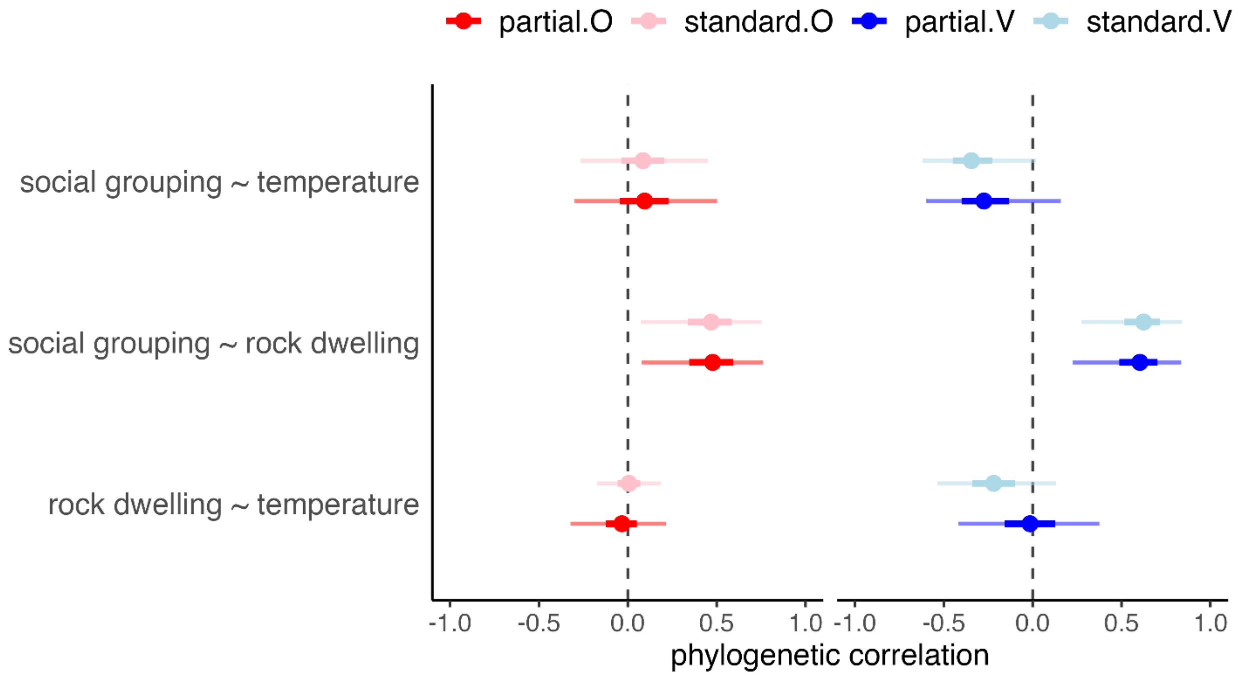


Figure S8. Standard and partial phylogenetic correlation estimates from MR-PMM estimating phylogenetic covariances for oviparous (O) and viviparous (V) species separately. Points represent posterior means, with heavy and light whiskers indicating the 0.5 and 0.95 credible intervals for each estimate. Data shown in table S8.
